# DNA Methylation Reprogramming during Sex Determination and Transition in Zebrafish

**DOI:** 10.1101/2020.09.17.301580

**Authors:** Xinxin Wang, Xin Ma, Gaobo Wei, Weirui Ma, Zhen Zhang, Xuepeng Chen, Lei Gao, Zhenbo Liu, Yue Yuan, Lizhi Yi, Jun Wang, Toshinobu Tokumoto, Junjiu Huang, Dahua Chen, Jian Zhang, Jiang Liu

**Affiliations:** CAS Key Laboratory of Genome Sciences and Information, Beijing Institute of Genomics, Chinese Academy of Sciences, Beijing 100101, China; University of Chinese Academy of Sciences, Beijing 100049, China; State Key Laboratory of Membrane Biology, Institute of Zoology, Chinese Academy of Sciences, Beijing 100101, China; State Key Laboratory of Molecular Developmental Biology, Institute of Genetics and Developmental Biology, Chinese Academy of Sciences, Beijing 100101, China; Key Laboratory of Reproductive Medicine of Guangdong Province, the First Affiliated Hospital and School of Life Sciences, Sun Yat-sen University, Guangzhou 510275, China; Integrated Bioscience Section, Graduate School of Science and Technology, National University Corporation Shizuoka University, Shizuoka 422-8529, Japan; Center for Excellence in Animal Evolution and Genetics, Chinese Academy of Sciences, Kunming 650223, China; Center for Life Sciences, School of Life Sciences, State Key Laboratory for Conservation and Utilization of Bio-Resources in Yunnan, Yunnan University, Kunming 650091, China

**Author notes:** These authors contributed equally to this work (Xinxin Wang, Xin Ma, Gaobo Wei). Correspondence to: Jiang Liu; Jian Zhang; Dahua Chen.

**Keywords:** Germ cell, DNA methylation, Sex determination, Sex transition

## Abstract

It is a mystery about sex determination and sexual plasticity in species without sex chromosomes. DNA methylation is a prevalent epigenetic modification in vertebrates, which has been shown to involve in the regulation of gene expression and embryo development. However, it remains unclear about how DNA methylation regulates sexual development. To elucidate it, we used zebrafish to investigate DNA methylation reprogramming during juvenile germ cell development and adult female-to-male sex transition. We revealed that primordial germ cells (PGCs) undergo significant DNA methylation reprogramming during germline development and set to an oocyte/ovary-like pattern at 9 days post fertilization (9 dpf). When blocking DNMTs activity in juveniles after 9 dpf, the zebrafish preferably develops into females. We also show that *Tet3* involves in PGC development. Notably, we find that DNA methylome reprogramming during adult zebrafish sex transition is similar to the reprogramming during the sex differentiation from 9 dpf PGCs to sperm. Furthermore, inhibiting DNMTs activity can prevent the female-to-male sex transition, suggesting that methylation reprogramming is required for zebrafish sex transition. In summary, DNA methylation plays important roles in zebrafish germline development and sexual plasticity.

## Introduction

Sex determination in animals has been one of the most fantastic mysteries in biology. In some species sex is determined by sex chromosomes, while in some other species no sex chromosomes can be found in their genomes [1]. It is rarely understood about the mechanism of sex determination in these species. Even more interestingly, some adult animals can change their sex in nature conditions. For example, sexual fate in *sea goldie* can be regulated by the presence or absence of male fish within the population. When a dominant male dies in one group, one of the remaining females in this group will transform into a functional male [2, 3].

Many fishes, such as domesticated laboratory zebrafish strains, do not have sex chromosomes [4]. Prior to the sex determination stage during zebrafish germline development, juvenile zebrafish have an ovary-like gonad which contains early stage oocytes [5]. If these early stage oocytes continue to mature, zebrafish will develop into females. On the contrary, if these oocytes start to degenerate, then “juvenile ovary to testis” transformation can be observed, and zebrafish will develop into males [6–8]. It has been found that zebrafish sexual fate can be affected by environmental factors, such as temperature and endocrine hormones [9–11]. Additionally, some studies have also reported that the germ cell number can influence sex determination in zebrafish [12]. In their natural growing environment, zebrafish sex transition can be observed. In the laboratory, aromatase inhibitor can induce female zebrafish to transform into male after sexual maturity [13]. However, the molecular mechanism about sexual plasticity in adults has not been largely elusive.

DNA methylation plays important roles in gene expression and development [14–18]. Two waves of DNA methylation reprogramming occur during mammalian early embryogenesis and primordial germ cell (PGC) development [19–24]. In mouse, abrogating DNA demethylation in primordial germ cells (PGCs) can lead to infertility [25], indicating the importance of DNA methylation programming during germ cell development. In zebrafish, the DNA methylation reprogramming during early embryogenesis has been investigated, showing that sperm but not oocyte DNA methylome is inherited by the offspring [26, 27]. Nevertheless, the role of DNA methylation during zebrafish sex differentiation and transition is largely unknown.

To address these issues, we systematically analyzed DNA methylome and transcriptome during zebrafish germline development and female-to-male transition. Our study provides the mechanism insights in understanding zebrafish sexual development.

## Results

### Mapping the global DNA methylome during zebrafish germ cell development

In order to investigate the dynamics of DNA methylome during zebrafish germline development, we collected the samples of 9 representative stages during PGC development and germ cell differentiation. We used a transgenic kop-gfp-nos1-3’UTR strain to collect early PGCs [28], including 4 hours post fertilization (4 hpf) PGCs at specification stage, 6 hpf PGCs at migration stage, and 24/36 hpf PGCs arriving at genital ridge (Figure 1A). kop GFP expression cannot be detected in later germline development stages, but vasa can mark later germ cells [29–31]. Thus, we generated *vasa::egfp* transgenic strain to trace later germ cells. We collected PGCs at 4 days post fertilization (4 dpf), 9 dpf, 17 dpf from juvenile gonads, and germ cells (stage-1B oocytes [32] with 30-50 um diameter) from 35 dpf female zebrafish (denoted as 35 dF germ cells) and germ cells (GFP positive cells with less than 10 um diameter) from 35 dpf male zebrafish (denoted as 35 dM germ cells) (Figure 1A and S1A). Zebrafish starts sex differentiation at 17 dpf [33]. In this study, germ cells from 4 hpf to 17 dpf stages are uniformly described as “PGCs”, and 35 dF and 35 dM germ cells are called as female germ cells (FGCs) and male germ cells (MGCs) respectively. Micromanipulation and fluorescence-activated cell sorting (FACS) methods were employed to isolate the germ cells (Figure S1B; see Methods). Highly purified germ cells were used to prepare DNA methylation libraries by low-input WGBS method [34] with at least two biological replicates (Table S1). CpGs covered by at least 3 reads were used for subsequent analysis. Our data show that CpG density is anti-correlated with DNA methylation level across all stages (Figure S2A). Most of genomic elements are hypermethylated except for 5’UTR and promoter CpG islands (CGIs) (Figure S2B). While, half of lncRNAs [35] show middle or low methylation levels, which is different from tRNA and rRNA (Figure S2C).

**Figure 1.**
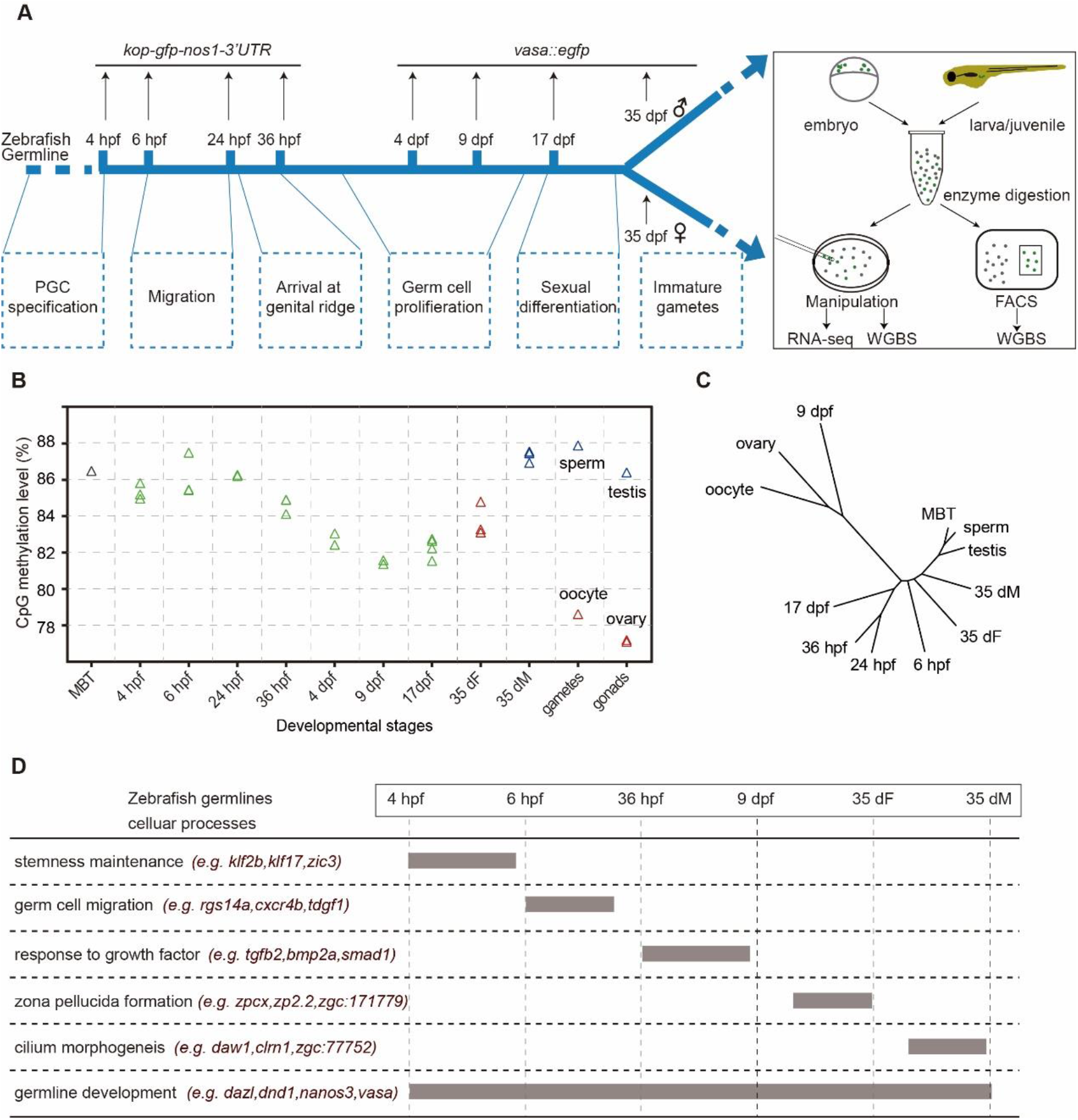
Dynamics of DNA methylation during zebrafish germline development. **A.** The schematic diagram for zebrafish germline developmental stages in this study. Two transgenic strains were used to track the development of germ cells in zebrafish. Dotted boxes indicate the major developmental processes of germ cells at different stages. **B.** Global DNA methylation levels during zebrafish germ cell development. DNA methylation data of zebrafish sperm, oocyte, MBT and testis stages are from public data (GSE44075). Green triangle means the PGC samples, red triangle means female samples, blue triangle means male samples, grey triangle means embryonic samples. **C.** A dendrogram showing the clustering of DNA methylomes during zebrafish germ cell development (500 bp bin for each unit). **D.** A brief summary of key gene expressions and key biological processes during zebrafish germline development.

Next, we plotted average methylation levels of PGCs at all examined stages. The global methylation levels do not substantially change for the early PGCs from 4 hpf to 24 hpf which are similar to that of midblastula (MBT) embryos (Figure 1B). This observation is similar to the results from the previous study [36]. Intriguingly, our data further show that after PGCs arrive at germinal ridge, the average methylation levels (MLs) of PGCs at 36 hpf start to decrease and reach the lowest point at 9 dpf (ML=81%) which is comparable to that of oocytes (ML=79%) and ovaries (ML=77%). The hierarchical clustering analysis of DNA methylomes also shows that 9 dpf PGCs, oocytes and ovaries cluster together, separated from the other stages (Figure 1C). This is different from previous study reporting that PGCs migrate into gonads maintain the stable methylome pattern [37]. Furthermore, the methylation level of 35 dM increases dramatically to a similar level of sperm. On the contrary, the methylation level of 35 dF is lower than that of 35 dM (Figure 1B). In summary, a unique reprogramming of DNA methylome occurs during zebrafish germline development, which is distinct from that in mammals [38].

### Transcription landscape in zebrafish germ cells

Besides DNA methylome, we also performed RNA-seq in zebrafish germ cells (Table S2). Firstly, we identified the stage-specific expressed genes during germline development (Figure S3A and Table S3). Gene ontology (GO) enrichment analysis shows that 6 hpf specifically expressed genes are enriched in the categories of somitogenesis and germ cell migration (Figure S3A), such as *G-protein signaling 14a* (*rgs14a*) (Figure S3B). It is consistent with the fact that germ cells at 6 hpf undergo the process of migration. Our data also show that pluripotency genes such as *zic3* and *klf17* are especially expressed at the stages of 4 hpf and 6 hpf (Figure 1D and S3B), suggesting that PGCs at 4 hpf and 6 hpf show more stemness. A significant number of genes are expressed in both 35dF and 35dM, but many genes are only highly expressed in 35dF or 35dM respectively. Further GO analyses show that 35 dF specifically expressed genes are enriched in the processes of egg coat formation and binding of sperm to zona pellucida (Figure S3A), such as *zp2.2* (Figure 1D and S3B), reflecting the character of oocytes. Notably, *zygote arrest 1-like* (*zar1l)*, *pou5f3* and *carbonic anhydrase 15b* (*ca15b)* are only expressed in 35 dF but not in 35 dM. These genes may function specifically in female germ cells (Figure S3B). The stage of 35dM specifically expressed genes are enriched in cilium assembly (e.g., *daw1*) (Figure 1D; Figure S3A and B), which is in line with the fact that the flagellum of sperm is a modified cilium. In addition, germ cell genes, such as *piwil* and *tdrd* family genes, show limited expression in early PGCs but highly express at the later stages Supplementary information, Figure S3B. However, genes like *vasa* and *dead end 1(dnd1)*, keep highly expressed throughout the whole examined development period (Figure 1D). Taken together, gene expression can reflect the general cellular processes of zebrafish germline development (Figure 1D).

### The role of methylation reprogramming during germ cell development

Promoters represent a vital regulatory level for transcription initiation [39]. DNA methylation in promoters can regulate gene expression [40]. Therefore, we focused on the DNA methylation dynamics at promoter regions. We performed clustering analysis based on differentially methylated promoters (DMPs) (Figure 2A). Our data show that the stage-specific hypomethylated promoters at the 9 dpf stage compared to the other developing stage are also hypomethylated in oocytes/ovaries, and the stage-specific hypermethylated promoters at the 9 dpf stage are hypermethylated in oocytes/ovaries as well. These results indicate that the DNA methylation pattern of 9 dpf PGCs at promoters is similar to that of the oocytes/ovaries.

**Figure 2.**
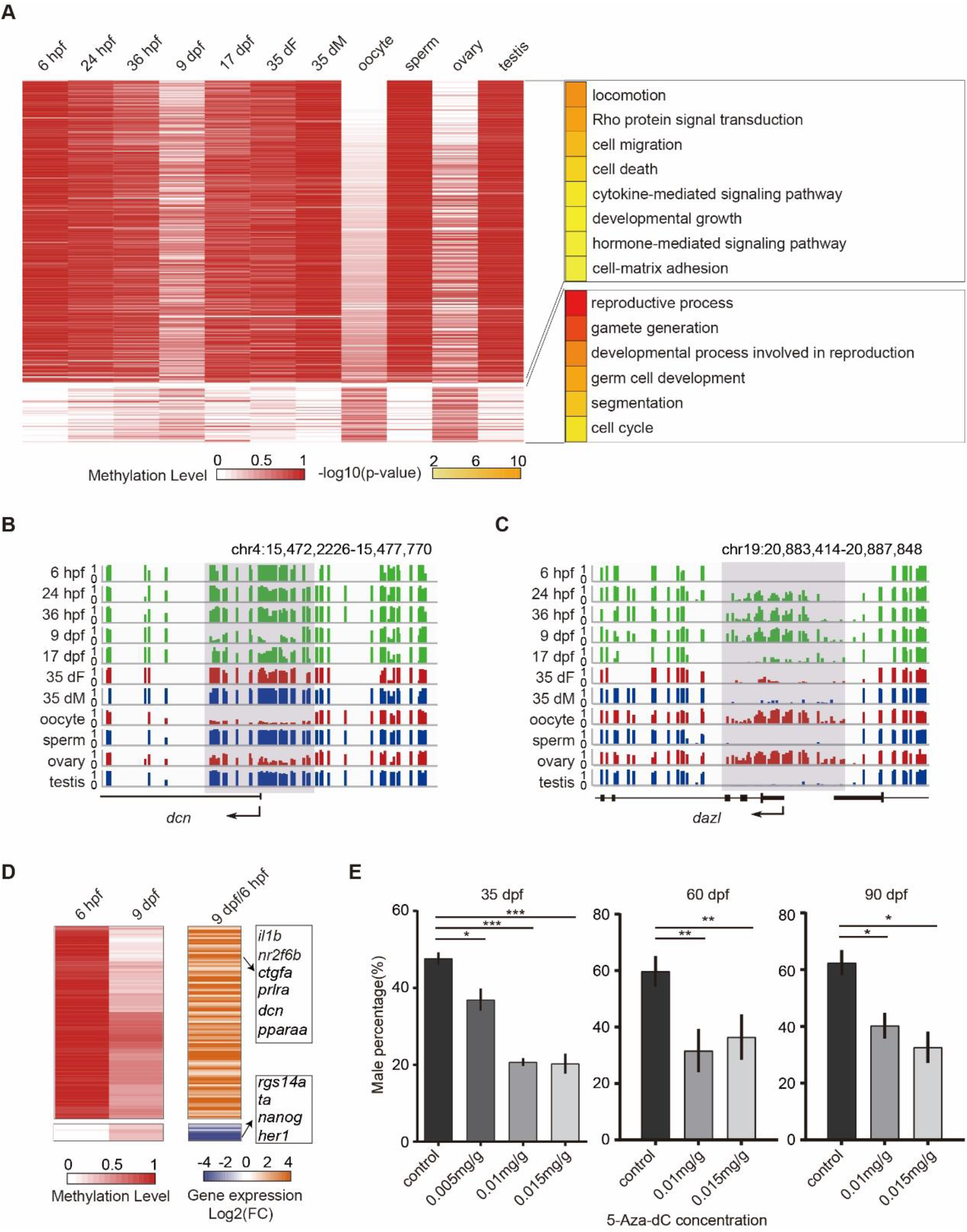
Methylome reprogramming during zebrafish germ cell development. **A.** DNA methylation level heatmap for differentially methylated promoters (DMPs). These DMPs are classified into groups by Partitioning Around Medoid (PAM) method. Corresponding GO biological process terms are shown in the right panel and the color represents the enrichment stastical significance. Promoters (Transcription Start Site, TSS±300 bp; n=385). **B.** Snapshots for DNA methylation at 9 dpf-specific hypomethylated promoter gene *dcn* (cytokine-mediated signal pathway related gene). Dynamic regions around promoters are highlighted in grey. DNA methylation level from 0 to 1. Each vertical line represents one CpG site. **C.** Snapshot for methylation at 9 dpf-specific hypermethylated promoter gene *dazl* (germline gene). **D.** Heatmaps of DNA methylation level (left) and gene expression fold change (right) for the genes with DMPs in 6 hpf and 9 dpf. Only genes with |log2 (fold change)|>1 are included. **E.** Sex ratio within zebrafish population after treatment with different 5-Aza-dC concentration, including 0.005mg/g (control n=124; 35 dpf n=47; 60 dpf n=121; 90 dpf n=112), 0.01mg/g (control n=86; 60 dpf n=90; 90 dpf n=82), 0.015mg/g (control n=60; 60 dpf n=64; 90 dpf n=56). The error bar is mean±s.e.m., Statistical significance were calculated by one-way ANOVA.

Further, GO enrichment analysis shows that genes with hypomethylated promoters in oocytes and 9 dpf PGCs are enriched in developmental growth, hormone mediated signal pathway and cell motion (Figure 2A and Table S4), such as *myoc* (Figure S4A), and *dcn* (Figure 2B) which regulates cell adhesion and migration by binding to ECM molecules [41]. The result is consistent with the observation that the PGCs migrate and then start to proliferate in gonads between 6 hpf and 9 dpf. Our data show that stage-specific hypermethylated promoters in 9 dpf PGCs are enriched in the reproductive process, germ cell development (Figure 2A and Table S4). These promoters become hypomethylated states at 17 dpf, 35 dF and 35 dM. The representative examples of germ cell development related genes *dazl*, *tdrd1* and *ta* respectively are shown in figures (Figure 2C; Figure S4B and C). We speculate that these 9 dpf hypermethylated-promoter genes could help PGCs avoid the precocious differentiation.

To examine the potential regulation of promoter methylation on gene expression during zebrafish PGC development, we also analyzed the relationship of differential gene expression and promoter methylation between 6 hpf and 9 dpf PGCs. Our results illustrate that gene expression usually increases when the promoter methylation level decreases, including cell migration, growth and hormone-mediated genes (e.g., *prlra*) (Figure 2D). By contrast, gene expression usually decreases when the promoter methylation level increases, including pluripotency gene (e.g., *nanog*) (Figure 2D). These results suggest that the dynamic of DNA methylation from 6 hpf to 9 dpf can inhibit the pluripotency capability of PGCs and facilitate PGCs migration and proliferation.

At 35 dpf, zebrafish has determined its sex. Gonads of female zebrafish at 35 dpf contain immature oocytes (Figure S1A) and show expression of oocyte-related genes (Figure S3B). However, there are a large amount of DMPs between 35 dF FGCs and oocytes, indicating that DNA methylome of 35dF FGCs are different from that of oocytes (Figure 2A). In contrast, the methylome pattern of 35 dM MGCs is quite similar to that of sperm (Figure 2A). These results suggest that the DNA methylome of sperm is nearly established as early as 35 dpf, whereas dramatic DNA methylation reprogramming is necessary for oocytes maturation after 35 dpf.

To further elucidate the role of DNA methylation on sex determination, we used 5-Aza-2’-deoxycytidine (inhibitors of DNA methyltransferases (DNMTs)) to block DNA methylation during zebrafish germ cell development. We utilized three dose-response 5-Aza-2’-deoxycytidine (5-Aza-dC: 0.005mg/g, 0.01mg/g, 0.015mg/g) to treat *vasa::egfp* transgenic strain at 9 dpf and raised the treated juvenile fish till 35 dpf, 60 dpf and 90 dpf. Then we checked the male/female ratio in each group (see Methods). The male sex ratio in treated group from 35 dpf to 90 dpf is lower than that in the control (Figure 2E and Table S5), indicating that fish treated with DNA methylation inhibitor are prefer to develop into females, which is consistent with recent study [42]. Therefore, these results suggested that DNA methylation plays a role during sex determination.

### *Tet3* involves in PGC development

Our data show there are more than ten thousand of demethylated differentially methylated regions (DMRs) between 36 hpf stage and 9 dpf stage (Figure S5A). *Tet* proteins were proposed to be involved in the DNA demethylation in mammalian embryos [43, 44] and zebrafish phylotypic-stage embryos [45]. Our data shows that *tet1* and *tet3* but not *tet2* were highly expressed in PGCs at the 36 hpf and 9 dpf stages (Figure 3A). To better define the role of *tet* during germ cell development, we used the morpholino (MO) knock down approach to target *tet1* and *tet3* in *vasa::egfp* transgenic embryos and validate by western blot (Figure S5B). The majority embryos injected with double morpholino (*tet1* MO + *tet3* MO) showed developmental deformity and lethality (Figure S5C). *Tet1* MO embryos did not show abnormal phenotypes. By contrast, *tet3* MO individuals showed much weaker fluorescence signal in gonads at 5 dpf and 9 dpf compared to the control (Figure S5D). Previous studies have reported that the germ cell number can affect sex determination in zebrafish [12]. Our results show that the number of PGCs in *tet3* MO juveniles is significantly reduced (Figure S5E).

**Figure 3.**
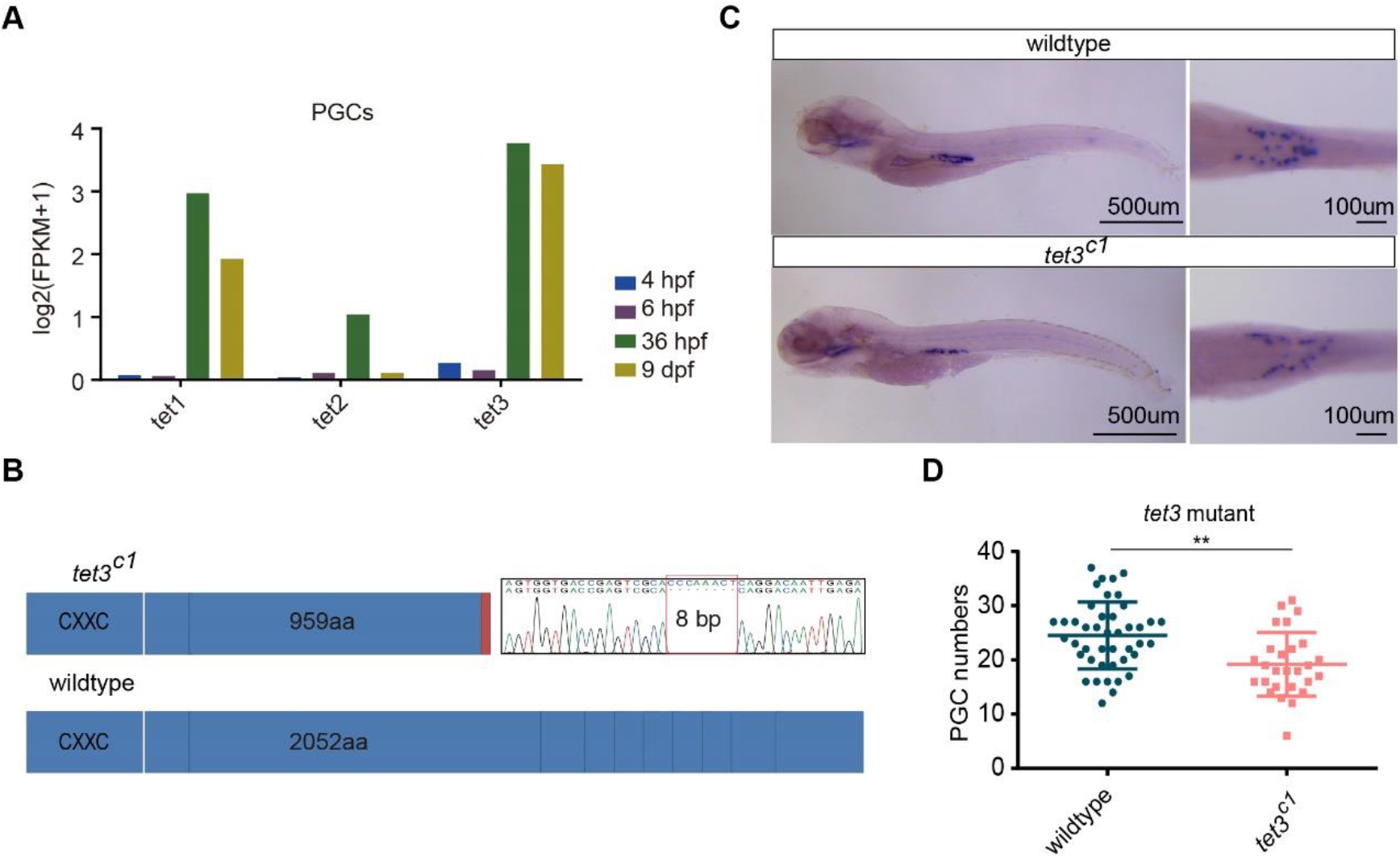
The function of *tet3*^*c1*^ mutant during zebrafish germ cell development. **A.** Histogram of the gene expression levels of *tet1/tet2/tet3* in PGCs. Gene expression level is averaged from biological replicates. **B.** Schematic diagram shows CRISPR/Cas9 editing in zebrafish *tet3* gene, which delete 8 bps in exon3. **C.** Expression of *vasa* mRNA in situ hybridization for wild type and *tet3*^*c1*^ mutant fish at the 5 dpf stage. **D.** The PGC number of 5 dpf was counted under stereoscopic microscope (wild type n=43; tet3 n=27). The error bar represents mean±s.d.. The p-value was calculated by t test. * means p < 0.05. ** means p < 0.01, *** means p < 0.001.

Next, we introduced mutations in the zebrafish *tet3* gene by using CRISPR/Cas9 method. The *tet3*^*c1*^ allele is a mutation allele with an 8-bp deletion in exon3. This deletion leads to a premature stop codon in exon3 and the loss of catalytic domain in C terminal (Figure 3B). Then, we counted the number of PGCs in *tet3*^*c1*^ mutants at 5 dpf under stereoscopic microscope by in situ hybridization with *vasa* probes (Figure 3C and D). We observed that *tet3*^*c1*^ fish contain less *vasa*-positive cells compared to the control, indicating that interrupting *tet3* gene can reduce germ cell numbers in zebrafish. Hence, the *Tet3* involves in the regulation of PGC development.

### Female-to-male sex transition in adult zebrafish

Inhibiting aromatase activity by aromatase inhibitor can revert adult zebrafish female-to-male [13]. This female to male sex transition process provides us a chance to explore the underlying molecular mechanisms for adult sexual plasticity in the vertebrate. Thus, we used aromatase inhibitor aromasin (Exemestane) to induce female-to-male sex reversal in *vasa::egfp* zebrafish (Figure 4A and B; See Methods). Our results show that the EGFP fluorescence intensity within adult gonads is gradually decreased after aromasin treatment, suggesting that the female germ cells start degeneration (Figure 4B). Histological staining also shows that the number of female germ cells is gradually reduced (Figure 4B). After aromasin treatment for three months, the gonad exhibits an intermediate transition state containing both degenerated oocytes and emerged male germ cells. We then stopped the aromasin treatment and found that the treated female fish would develop a testis-like gonad in the next one or two months (Figure 4B). We further mated the sex reversed fish with wild type females. Our result shows that the sperm from sex reversal fish is able to successfully fertilize and produce normal embryos (Figure S6A).

**Figure 4.**
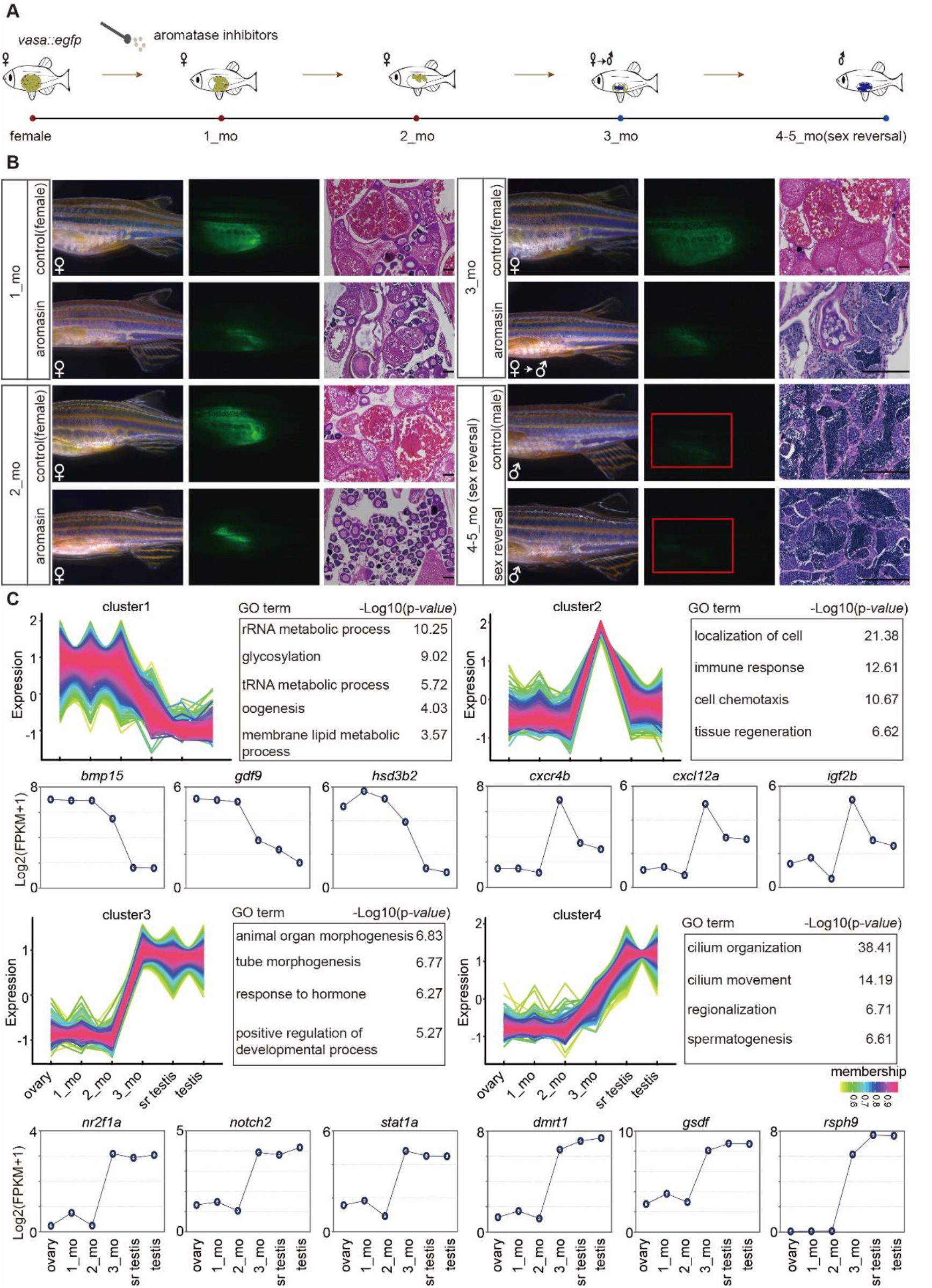
Female-to-male sex transition in the adult zebrafish. **A.** The schematic diagram for zebrafish sex reversal samples in this study. *vasa::egfp* transgenic strain was used to induce sex reversal by aromasin treatment. The samples in this figure include wild type ovary, gonads from zebrafish with one month (1_mo), two month (2_mo), and three month (3_mo) drug treatment, sex reversal testis and wild type testis. **B.** Observation of gonad morphology change in *Tg* (*vasa::egfp*) wild type female and post aromasin treatment fish per month. The image of gonads under bright filed, GFP fluorescence and HE staining. The scale bar indicates 100 μm. **C.** Time-course clustering analysis of gene expression patterns across zebrafish sex reversal process. In the four distinct expression groups, the upper left panel of each group shows the gene expression pattern, the upper right panel shows the GO enrichment results, and the representative genes expression dynamics is shown in the lower panel.

We then performed RNA sequencing to investigate the dynamics of gene expression during female-to-male sex transition (Table S2). By the time-course gene expression clustering analysis (see Methods), we identified four dynamic gene clusters (Figure 4C and Table S6). Our data show that the expression level for female sexual development genes (cluster1, e.g., *bone morphogenetic protein 15* (*bmp15)* and *hsd3b2*) are down-regulated during the sex transition process (Figure 4C). It is worth noting that 3_mo stage is intermediate transition stage (Figure 4B). Genes (cluster 2) which display uniquely high expression levels at this stage but low expression in the ovary and the sex reversed/ wild type testis were enriched in the tissue regeneration process (e.g., *igf2b*), immune response and localization of cell process (e.g., *cxcr4b and cxcl12a*) (Figure 4C). We also found genes which became highly expressed and kept high levels in the sex reversed/wild type testis (cluster3, e.g., *notch2*) were enriched in organ morphogenesis and hormone response (Figure 4C). In the end, male sexual development genes (cluster4, e.g., *doublesex and mab-3 related transcription factor 1* (*dmrt1)* and *gonadal somatic cell derived factor (gsdf)*) are up-regulated. In summary, during female-to-male transition, genes related to female sexual development stop expression at first, then genes related to tissue regeneration, cell migration, organ morphogenesis and hormone response start expression at the inter transition stage, and genes related to male and sperm generation start expression in the end.

### DNA methylome reprogramming during adult zebrafish female-to-male sex transition

Our results have shown that the DNA methylation is tightly associated with germ cell development in juvenile zebrafish (Figure 2). To explore the role of DNA methylation in adult zebrafish sex transition, we sequenced DNA methylomes of gonads during sex transition process, including wild type ovary/testis, gonads treated by aromasin for 1 month (1_mo), 2 months (2_mo), and 3 months (3_mo) and fully sex reversal testis (sr testis) with at least two independent biological replicates (Figure 4A; Figure S6B and Table S1). Our data show that the global DNA methylation level has a significant change during female-to-male sex transitions (Figure 5A), and the methylation level of 3_mo gonads (ML= 83%) is between that of wild type ovary (ML=77%) and testis (ML=86%) (Figure 5A and B; Figure S6C). When the treated females sexually revert to males, the methylation level of gonads from sex reversal fish reached to a similar level of wild type testis (Figure 5A and B; Figure S6C). The results above indicate that sperm from sex reversal fish have same functions as wild type sperm (Figure S6A). Then we also sequenced DNA methylomes for sex reversal sperm and wild type sperm. Our data show that the DNA methylome of sperm from sex reversal fish is similar to that of sperm from wild type fish, but different from that of oocytes (Figure 5C). For examples, *dnmt6* is low-methylated in oocyte, but becomes high-methylated in sex reversal sperm, which is similar to the state in wild type sperm (Figure S6D) and *sycp3l* is high-methylated in oocyte, but show low-methylated in sex reversal sperm (Figure S6E).

**Figure 5.**
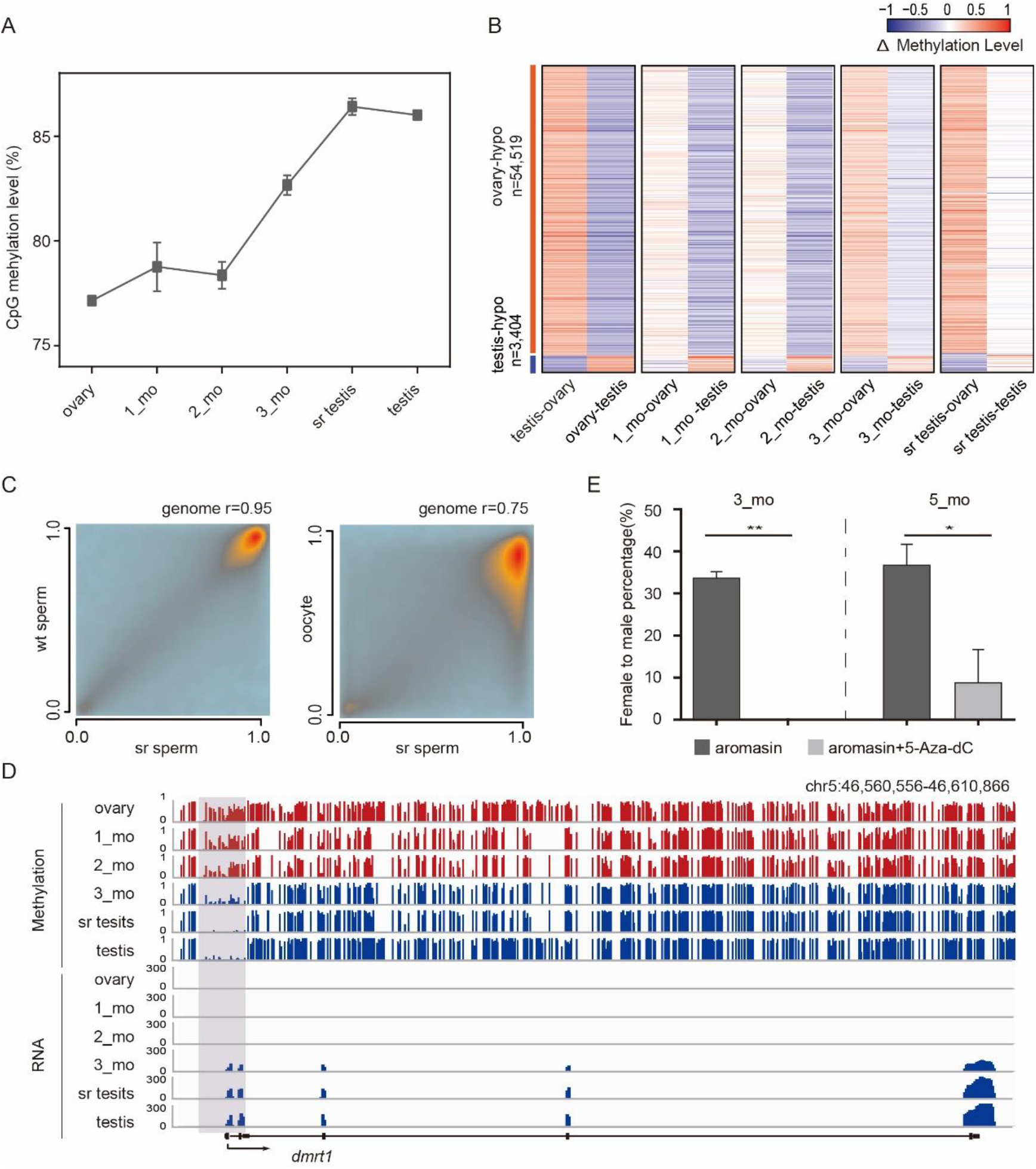
DNA methylation reprogramming during zebrafish female-to-male transition. **A.** The global DNA methylation reprogramming during zebrafish sex reversal. The methylation level is calculated as the average across all covered CpGs. The error bar represents mean±s.d.. The samples in this figure include wild type ovary, gonads from zebrafish with one month (1_mo), two month (2_mo), and three month (3_mo) drug treatment, sex reversal testis and wild type testis. **B.** The dynamic changes for DMRs between zebrafish ovary and testis during sex transition process. The △ methylation heatmap displays the difference of DNA methylation level between chosen two stages. **C.** The correlation analysis of CpG methylation level in sex reversal sperm (sr sperm), wild type sperm (wt sperm) and oocyte. Pearson correlation coefficients r is also shown in the figure. **D.** Snapshots of DNA methylation and gene expression tracks for sex determination gene *dmrt1* in ovary, 1_mo, 2_mo, 3_mo, sr testis and wt testis. **E.** Female sex ratio within populations after drug treatment for the 3_mo stage (aromasin n=36; aromasin+5-Aza-dC n=47) and 5_mo stage (sex reversed stage, aromasin n=36; aromasin+5-Aza-dC n=38). 3 independent experiments for each drug treatment group. The error bar is mean±s.e.m., the statistical significance was calculated by unpaired two-sided t test.

Our data illustrate that several well-known sex determination genes show differentially methylated at their promoters between ovaries and testis (Figure S7A and B; Table S7). Then, we investigated the relationship between transcription and methylation status for these known sex determination genes during sex transition. For example, a well-defined sex determination gene *dmrt1*, a key regulator of sex determination which is necessary for male sexual development in vertebrates [46–49], has a male-specific hypomethylated promoter and starts to be highly expressed from 3_mo onwards (Figure 5D). In addition, *cytochrome P450 superfamily member-cyp19a1a*, a critical gene in promoting ovary development in fishes [50], shows a low methylation level at the promoter region in ovaries, but gradually reprograms to a high methylation level at the later stages (Figure S7C).

We next compared the DNA methylation level dynamics between adult sex transition process and juvenile germline development in zebrafish. The dynamics of methylation level during female-to-male sex transition is very similar to that from 9 dpf germ cells to testis during normal development (Figure S7D and E). Perhaps, the methylome of female gonads similar to the 9 dpf-like bipotential DNA methylation pattern is the reason why the female gonad is able to transform into the male gonad.

Furthermore, we wanted to know the role of DNA methylome during zebrafish sex reversal process. We added 5-Aza-dC (0.035mg/g) to examine its effect on aromasin-induced female-to-male sex transition (see Methods). After aromasin treatment for three months, 31% of total female fish with only– aromasin-treatment start transiting to male as these fish have both ovary and testis, while none of females with both 5-Aza-dC and aromasin treatment start the transition. Furthermore, we raised the treated fish in the recirculation system till 5 months after the initial treatment began (fish sex reversed stage) with no more drug treatment and found that 33% of total females transited into males for the only-aromasin-treatment group, whereas merely 5% of females transited into males for both 5-Aza-dC and aromasin group (Figure 5E and Table S5).

In summary, our results indicate that blocking DNA methylation can prevent sex reversal in zebrafish, suggesting that DNA methylation dynamics is essential for sex reversal in adult zebrafish.

## Discussion

Genome-wide erasure of DNA methylation during the germline development has been documented in mammals [21–24, 38], but it was limitedly reported in zebrafish germ cells. A recent study generated low depths of DNA methylome of zebrafish PGCs after 24 hpf [37]. Low depths and coverage of the DNA methylomes block the understanding of germ cell development. They report that zebrafish did not experience significant reprogramming during PGC development. In contrast, we generated a relatively higher depths of DNA methylomes for zebrafish including early PGCs and late germ cells using modified library generation methods at single-base resolution [34]. Our results show that zebrafish PGCs undergo significant DNA methylation reprogramming during the germline development. More importantly, we further reveal that the methylome reset to an oocyte/ovary-like pattern at 9 dpf stage, and illustrate that zebrafish PGCs at 9 dpf have the bipotential to differentiate into male or female germ cells. It is excited for us to find that a similar methylation reprogramming during normal sexual development from 9 dpf PGC to male testis can be observed during aromasin-induced female ovary to male testis transition. Thus, we infer that the 9 dpf-like bipotential DNA methylation pattern in female gonads is the reason why the female gonad is able to transform into the male gonad. The role of DNA methylation in sex reversal had not been revealed previously. In this study, we inhibit the DNA methylation reprogramming after aromasin treatment, and demonstrate that DNA methylation reprogramming is required for the sex reversal in zebrafish.

During artificially induced zebrafish female-to-male sex transition, our results indicated that there exists DNA methylation dynamics for vital sex determination genes including dmrt1 and cyp19a1a, which shows a similar trend in blue head wrasses and half-smooth tongue sole[51, 52]. These results suggest that DNA methylation may have a conserved regulatory role in fish sex determination. Recent study report that H3K27me3 plays roles in turtle sex determination [53]. The interplay of DNA methylation and other epigenetic mechanisms for zebrafish sex determination is worthy of further study.

Taken together, our research provides molecular mechanism insights for understanding the mystery of sex reversal, and we revealed that the DNA methylome reprogramming play roles in zebrafish germline development and female-to-male sex transition.

## Competing interests

The authors declare no competing interests.

---

**Supplementary Figure S1.**
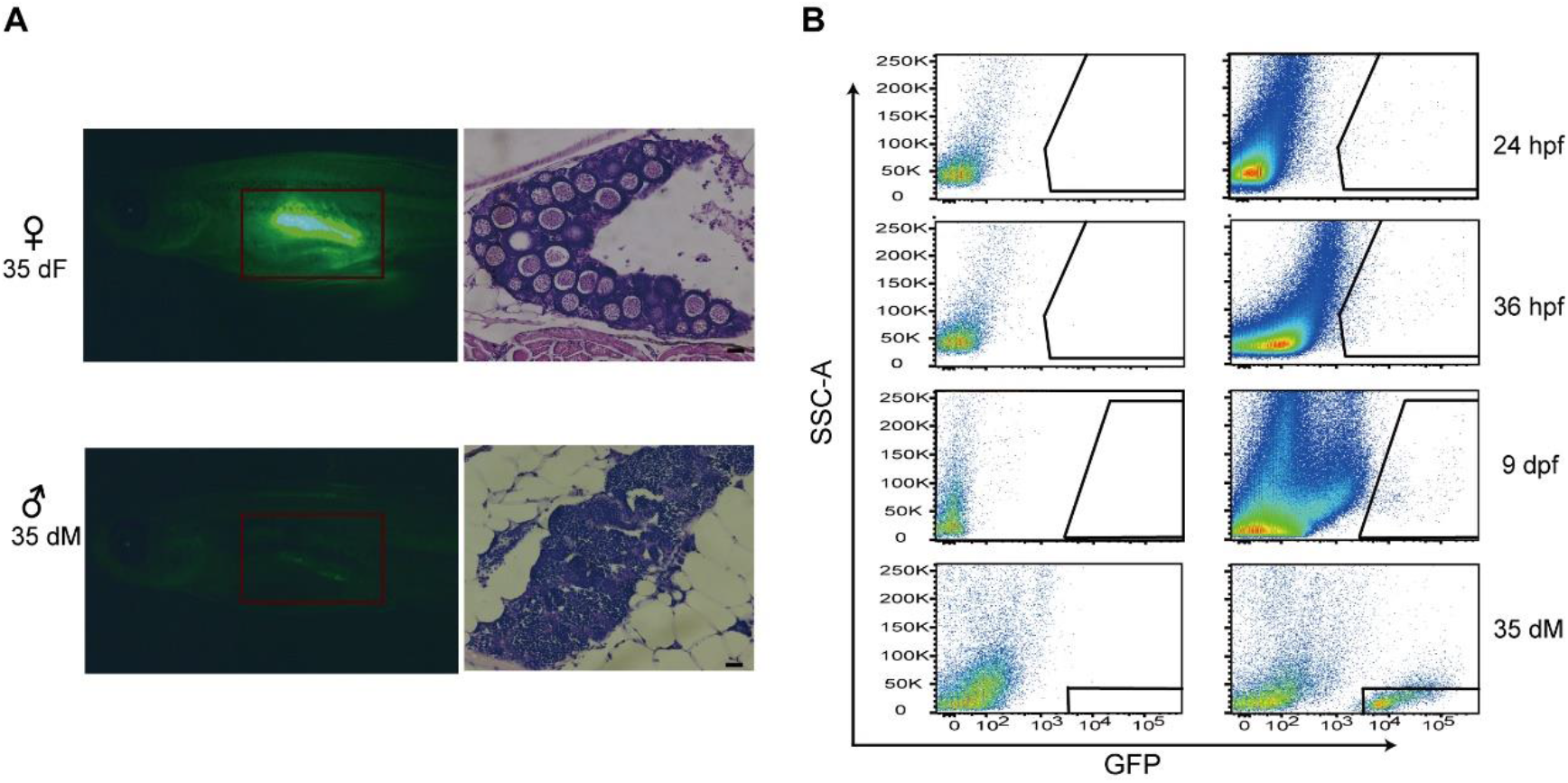
PGC and germ cell purification. **A.** Fluorescent photography and HE staining for immature ovary and immature testis at 35 dpf stage (approximately equal length of males and females around 1.7±0.1cm). The left panel represents the fluorescence intensity of ovary and testis at 35 dpf; the right panel represents the HE staining of ovary and testis at 35 dpf. The scale bar is 50 μm. **B.** Sorting GFP positive cells by FACS. Left panel shows GFP negative cells for control fishes. Right panel shows GFP positive cells at each germline time point for transgenic fishes.

**Supplementary Figure S2.**
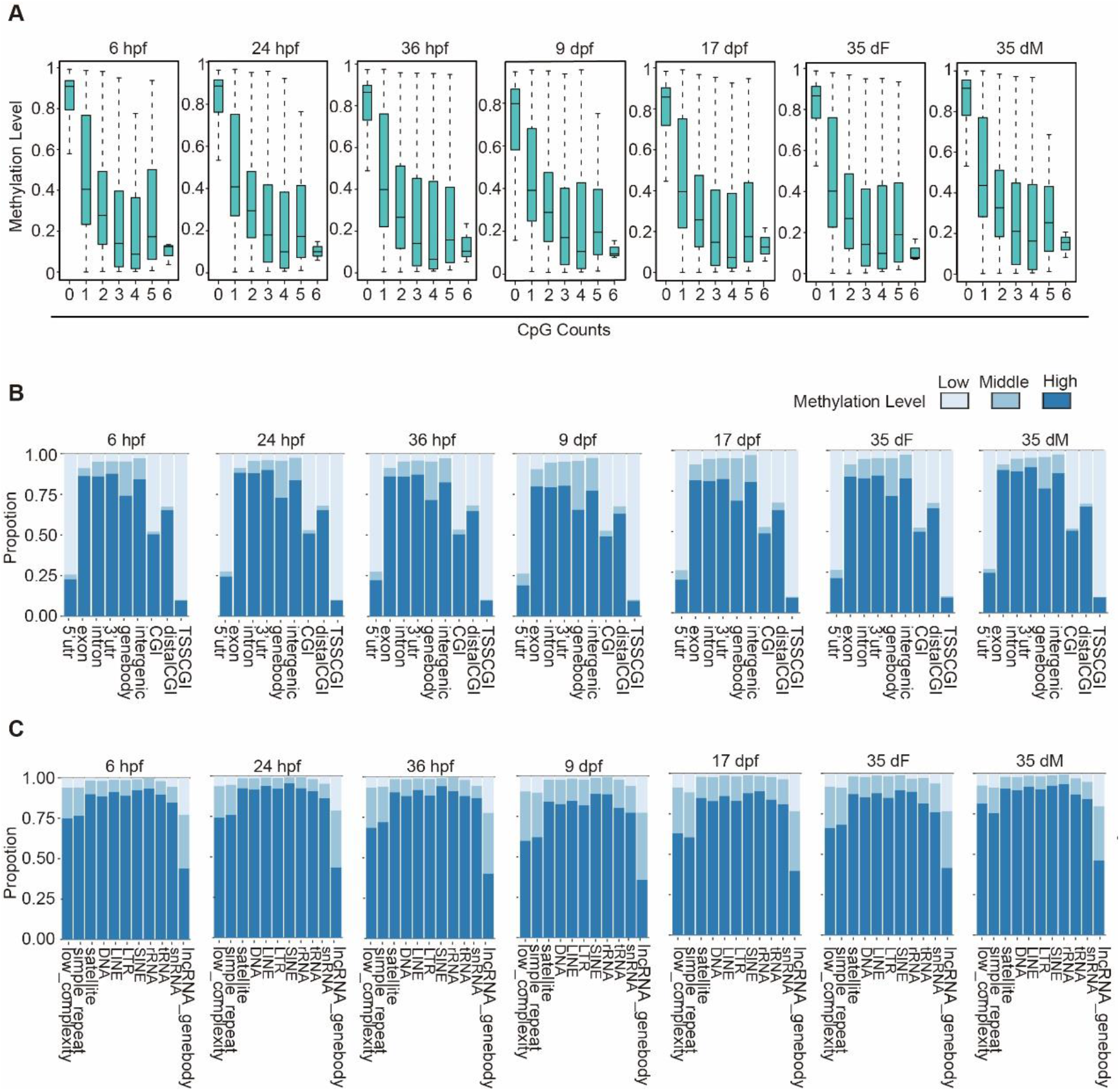
Global analysis of DNA methylation at different germline developmental stages in zebrafish. **A.** Boxplots show the correlation between CpG density and methylation level around TSS±2Kb regions. Boxes and whiskers represent the 25th/50th/75th percentiles and 1.5X the interquartile range, respectively. X axis means CpG counts. **B and C.** The Proportion of CpGs with high methylation level (ML ≥0.75), middle methylation level (0.25 < ML < 0.75) and low methylation level (ML ≤0.25) for indicated genomic features: 5’ untranslated regions (UTRs), exons, introns, 3’UTRs, gene body, intergenic regions and CpG islands (CGIs) are shown in **B** and repeat elements, noncoding RNAs are shown in **C**.

**Supplementary Figure S3.**
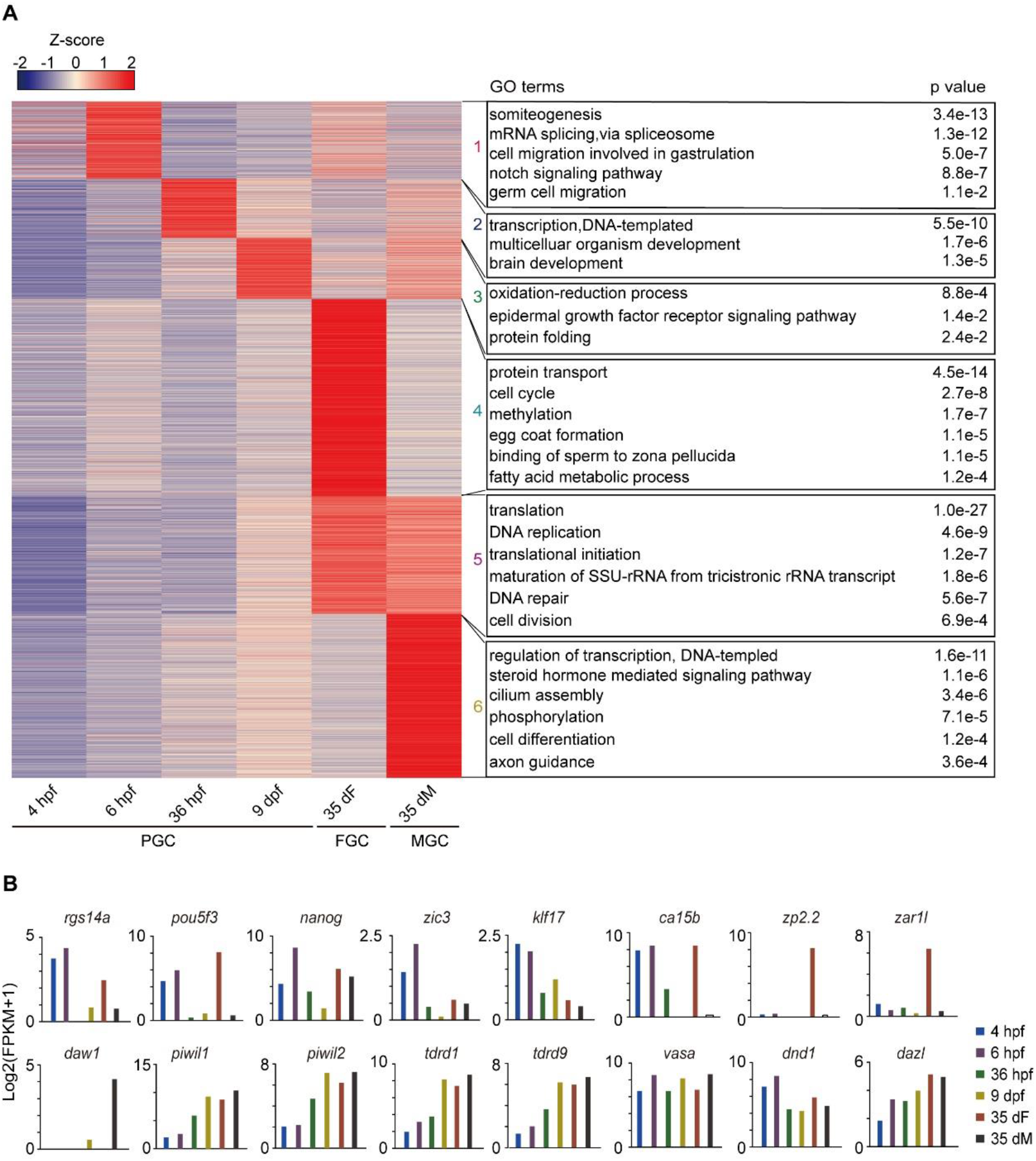
Transcription landscapes of zebrafish germ cells. **A.** Heatmap for stage specific gene expression from 4 hpf PGCs to 35 dpf germ cells. 6 clusters were identified by k-means method. 1253, 952, 989, 3191, 2717, 1902 genes for each cluster. The right panels show the corresponding cluster-enriched GO biological process terms by using DAVID. P-values are also shown. **B.** Barplots for the gene expression levels of germline-related genes at different stages. Gene expression levels are averaged from all biological replicates.

**Supplementary Figure S4.**
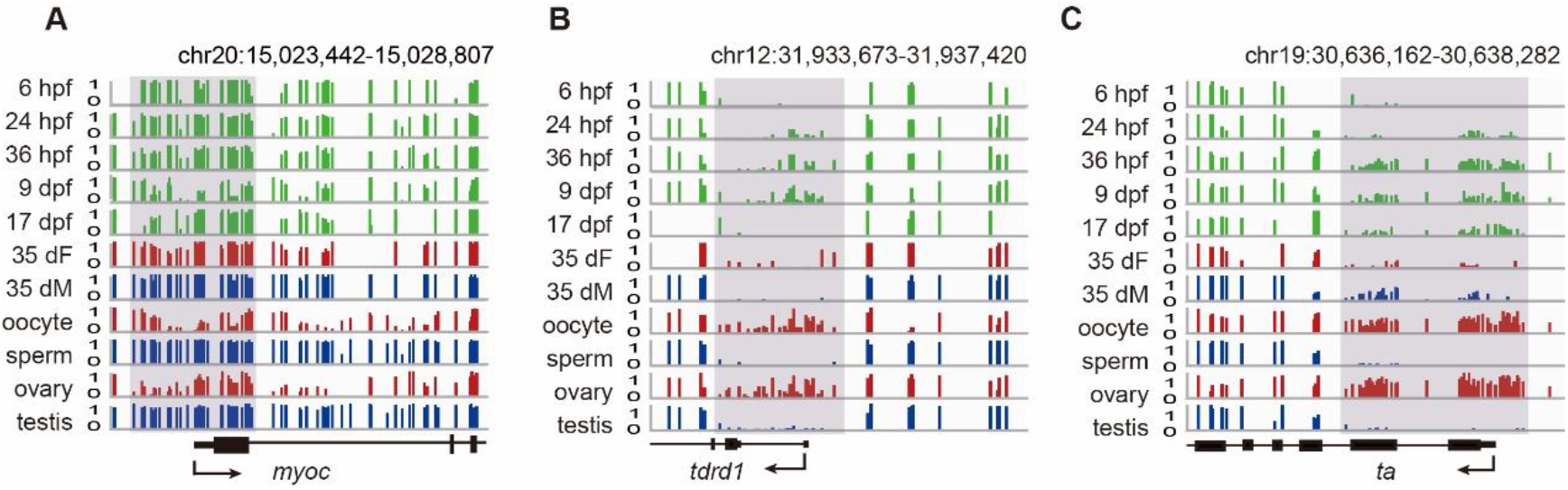
DNA methylation dynamics at promoter regions across zebrafish germline development. **A-C.** Snapshots for the DNA methylation of 9 dpf-specific hypomethylated promoter gene *myoc* **A** and 9 dpf-specific hypermethylated promoter gene *tdrd1* **B** and *ta* **C**. Dynamic regions around promoters are highlighted in grey. DNA methylation level from 0 to 1. Each vertical line represents one CpG site.

**Supplementary Figure S5.**
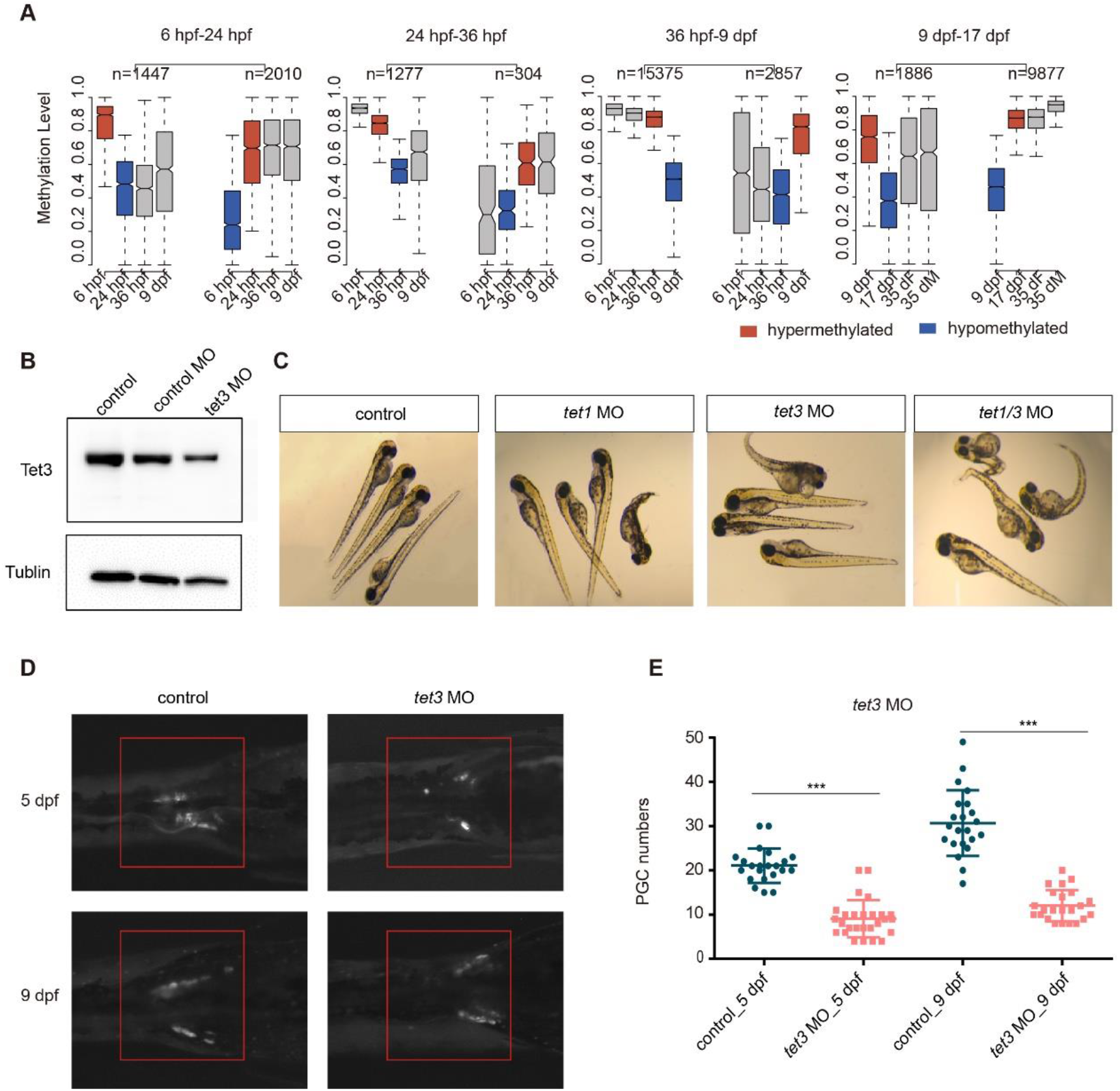
The function of *tet3* morpholino in zebrafish germ cell development. **A.** Boxplots of CpG methylation level in differentially methylated regions (DMRs) identified by stage adjacent pairwise comparisons. The hypermethylated stage is denoted as red box and the hypomethylated stage is denoted as blue box, grey boxes stand for other stages. Boxes and whiskers represent the 25th/50th/75th percentiles and 1.5X the interquartile range, respectively. n represents the number of DMRs. **B**. The western blot for control larvas, control MO larvas and *tet3* MO larvas at 3 dpf. Tubulin is the loading control for western blot. **C.** The images for control larvas, *tet1* MO larvas, *tet3* MO larvas and *tet1/3* MO larvas at 3 dpf under microscopy. **D.** The fluorescence intensity for wild type and *tet3* MO germ cells at 5 dpf and 9 dpf (red box). **E.** Dot plot showing changes in the PGC number at 5 dpf (control n=22; tet3 MO n=27) and 9 dpf (control n= 22 tet3 MO n=23), which was counted by the squash method. The error bar represents mean±s.d.. The p-value was calculated by t test. * means p < 0.05. ** means p < 0.01, *** means p < 0.001.

**Supplementary Figure S6.**
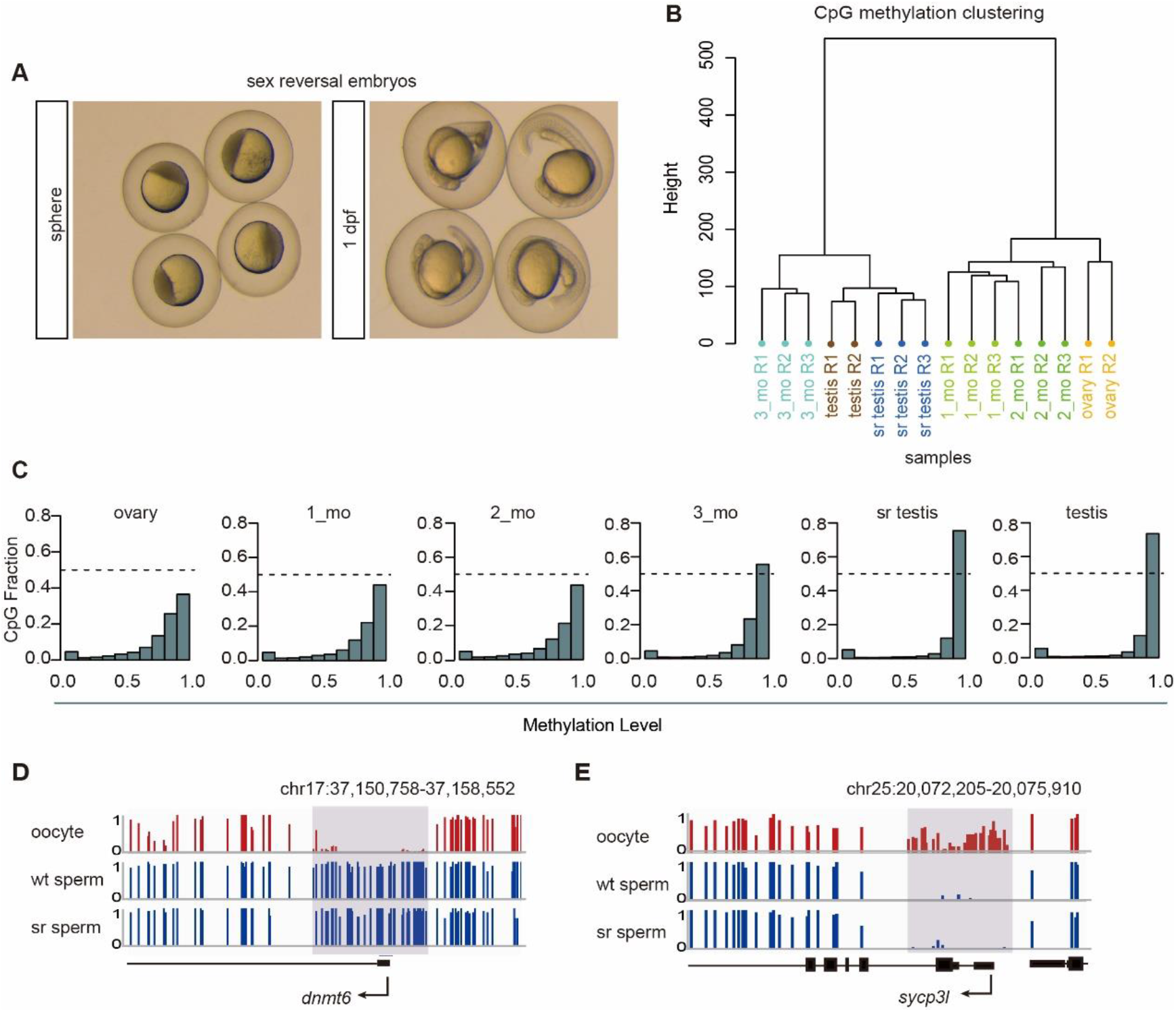
DNA methylation dynamics for zebrafish sex reversal samples. **A.** The pictures for embryos generated by sex reversal fish mating with wild type female. **B.** Hierarchical clustering analysis according to the DNA methylation pattern for the biological replicates of ovary, 1_mo, 2_mo, 3_mo, sr testis and testis. sr means sex reversal. **C.** Bimodal distribution of CpG methylation level across genome for zebrafish at the indicated stages: ovary, 1_mo, 2_mo and 3_mo drug treated gonad, sr testis and testis. **D** and **E.** Snapshots of DNA methylation tracks for genes with DMPs among oocyte, wt sperm and sr sperm, including *dnmt6* **D** and *sycp3l* **E**.

**Supplementary Figure S7.**
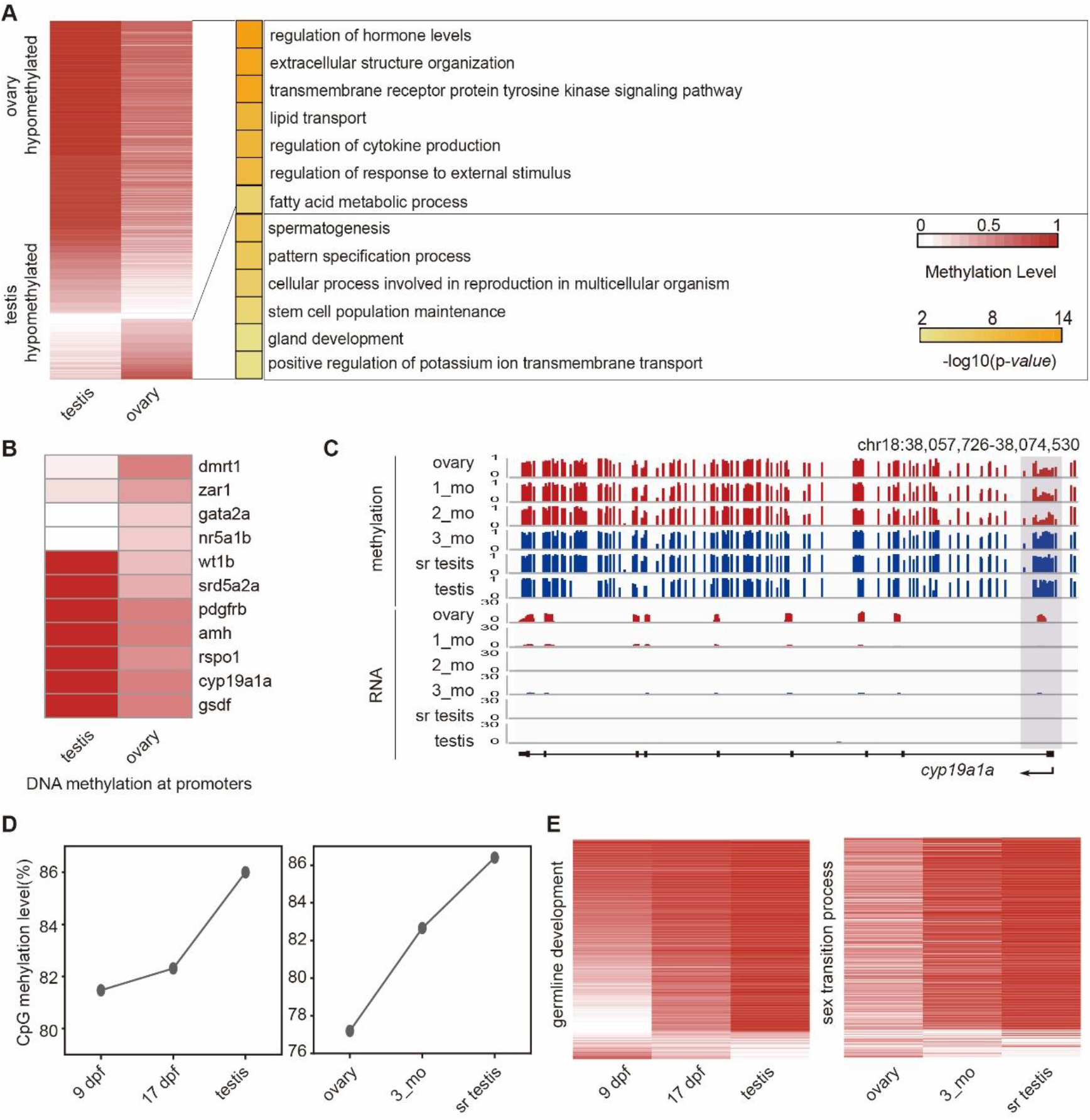
The relationship between DNA methylation and gene expression during zebrafish sex transition. **A.** DMPs between testis and ovary. GO terms for the testis-specific hypomethylated and ovary-specific hypomethylated DMPs are shown in the right panel. Promoters (TSS±1000 bp). **B.** Heatmap shows DMPs between ovary and testis for key sex determination genes. **C.** Snapshots of DNA methylation and gene expression tracks for sex determination gene *cyp19a1a* in ovary, 1_mo, 2_mo and 3_mo drug treated gonad, sr testis and testis. **D.** The comparison of DNA methylation reprogramming during sex transition process and germline development process in zebrafish. **E.** The heatmap shows the dynamic changes of DMPs between 9 dpf PGCs and testis during sex transition process (DMPs, n=574).

## SUPPLEMENTARY TABLES (tables separately in Excel format)

**Table 1.** DNA methylation data summary for zebrafish germline and sex reversal samples

**Table 2.** Transcriptome data summary for zebrafish germline and sex reversal samples

**Table 3.** Go terms for stage specific gene in zebrafish germ cells

**Table 4.** Go terms for differentially methylated promoters (DMPs) in zebrafish germline

**Table 5.** Sex ratio summary for drug treatment fish

**Table 6.** Go terms for time-course clustering genes during zebrafish sex transition

**Table 7.** Go terms for DMPs between testis and ovary

## Notes

### Competing Interest Statement

The authors have declared no competing interest.

### Summary of Updates

Adjust author order

