## supplementary method for "DNA Methylation Reprogramming during Sex Determination and Transition in Zebrafish"

### Materials and methods

Experimental model and subject details

All animal maintenance and experimental procedures were carried out according to the guidelines of the Institutional Animal Care and Use Committee (IACUC) of the Beijing Institute of Genomics, CAS, Beijing, China. All zebrafish used in this study were maintained at 28 °C with a light/dark cycle of 14/10 hours.

Generation of the transgenic zebrafish lines

The transgenic kop-gfp-nos1-3'UTR zebrafish line (AB strain based)was a kind gift from Jian Zhang Lab (Institute of Genetics and Developmental Biology, CAS, Beijing, China). Briefly, kop-gfp-nos1-3'UTR sequence was cloned into T2ASAd vector. The purified T2ASAd plasmid was injected into the cytoplasm of 1-cell stage zebrafish embryos with Tol2 transposase mRNA. Female fishes carrying the transgene were identified by screening of GFP expression in early PGCs.

The transgenic *vasa::egfp* zebrafish line (AB strain based) was a kind gift from Zhou Songyang Lab (Sun Yat-sen University, Guangzhou, China). Briefly, *Xba*I and *SfiI* restriction sites of pK4 vector[31] were used to release the *vasa::egfp* fragment, then the fragment was purified and injected into the cytoplasm of 1-cell stage zebrafish embryos. Injected embryos were screened for EGFP signals under a fluorescent microscope as founder candidates and cultured to sexual maturity. Female founder candidates were mated with wild type males and screened for the production of eggs with EGFP signals.

Sample collection and preparation of single cell suspension

For 4 hpf PGC and 6 hpf PGC collection, *kop-gfp-nos1-3'UTR* transgene zebrafish embryos were washed for two times with ice-cold 1X DPBS (Gibco), and chorions were carefully removed with sharp tweezers. 1 ml DPBS with 0.5% BSA was used to collected embryos, and gentle pipetting until all embryos were dissociated to single cells. All steps were operated on ice. For 24 hpf PGC and 36 hpf PGC collection, *kop-gfp-nos1-3'UTR* zebrafish embryos were placed on dish and depigmented by treating them with 1-Phenyl 2-thiourea (PTU, Sigma-Aldrich P7629). Chorions were removed with pronase (Roche) for 10 mins, and embryos were washed for two times with ice-cold DPBS. Embryo yolks were removed with 1 ml ice-cold deyorking buffer (55 mM NaCl, 1.8 mM KCl, 1.25 mM NaHCO3). Then 0.25% trypsin (Gibco) solution was added, and samples were incubated at 30°C for 15 mins followed by being washed in DPBS and centrifuged at 400 xg for 3 mins at 4°C. Then isolated single cells were re-suspended in DPBS containing 5% EDTA.

For 4 dpf and 9 dpf PGC collection, vasa::egfp zebrafish juvenile heads and tails were removed and body part containing gonads were dissected into D-hanks (Leagene) solution, then transferred into 0.25% trypsin and incubated at 30°C for 40 mins. Gentle pipetting was performed several times. The reaction was quenched with 10% fetal bovine serum (FBS), and individual cells were washed in DPBS and centrifuged at 400 xg for 3 mins at 4°C. The cell pellet was re-suspended in DPBS with 0.5% BSA.

For 17 dpf, 35 dpf female and 35 dpf male germ cells collection, gonads from individual zebrafish were isolated and separated from surrounding tissue in D-hanks solution with the help of a dissecting fluorescence microscope. Immature females and males were determined via GFP intensity of *vasa::egfp* zebrafish under a fluorescence microscope. Gonads exhibiting high and low EGFP fluorescence were classified as ovaries and testes, respectively [7]. The collected gonads were dissociated with 0.25% trypsin at 30°C for 10 mins. 10% FBS was added for termination of trypsin as described above. 35 dpf female germ cells (the stage-1B oocyte [32] with 30-50-um size were selected in this study) were centrifuged at 100 xg for 1mins at 4°C to avoid cell membrane breakdown.

For each sample, GFP positive cells were manually picked using micromanipulator system under the fluorescence microscope or were purified by FACS using FASAria cell sorter. For FACS, the cell suspension was filtered with 30 um Pre-Separation Filters (Miltenyi Biotec #130-041-407) to remove cell aggregates before sorting. GFP positive cells (less than 10 um diameter) at 35 dpf male fish which were potential spermatocyte, were sorted for further analysis. Cell purity was checked by fluorescence under microscopy.

Low-input bisulfate-seq library preparation for zebrafish germ cells

DNA methylation libraries of zebrafish germ cells were prepared with the low-input “one Tube” method as described previously [54]. 50-1000 cells lysed in 20 ul lysis buffer (20 mM Tris, 2 mM EDTA, 20 mM KCl, 1 mg/mL protease K) for 1.5 hours at 56°C, 30 mins at 75°C for inactivated. 30 ul nuclease free water and 0.5% spike-in un-methylated lambda DNA (Promega) were added into the lysate. DNA was sheared into 400 bp fragments by Covaris S2. Fragmented DNA was then concentrated to 30 ul and subjected to end repair by incubating with 5 ul end-repair enzyme mixture (3.5 ul T4 DNA ligase buffer (NEB), 0.35 ul 10 mM dNTP, 1.15 ul NEBNext End Repair Enzyme Mix) for 30 mins at 20°C followed by heat-inactivation for 30 mins at 75°C. After end-repair, 5 ul of dA-tailing mixture [0.5 ul T4 DNA ligase buffer, 1 ul Klenow exo- (NEB), 0.5 ul 100 mM dATP and 3 ul nuclease free water] was added to the tube and incubated for 30 mins at 37°C followed by heat-inactivation for 30 mins at 75°C. Finally, 10ul ligation mixture (1ul T4 DNA ligase buffer, 0.5 ul 100 mM ATP, 1.5 ul 5 mM cytosine methylated Illumina adapter, 2 ul T4 DNA ligase (NEB) and 5 ul nuclease free water) was added to the tube and incubated at 16°C overnight. 100 ng Carrier RNA was added into the tube. Bisulfite conversion reaction is performed with the EZ DNA Methylation-Gold Kit (Zymo Research) according to the manufacturer’s instructions. DNA was purified with column purification. The purified DNA was then amplified with 6 cycles PCR by using KAPA HiFi HotStart Uracil+ ReadyMix (KAPA). Amplified DNA was purified with 1 volume Ampure XP beads (Beckman Coulter A63881) to discard the short fragments and adapter-self ligations. 50 µl of XP beads was added to 50 µl of PCR product.

Samples were mixed by pipetting and incubated at room temperature for 10 mins. Keeping the beads on the magnet, samples were washed twice with 200 μl of 80% ethanol without mixing. Ethanol was then completely removed. Beads were left on the magnet for 5 mins to allow the remaining ethanol to evaporate. DNA was eluted by adding 20 μl of ddH2O, mixing by pipetting, and incubating for 5 mins. After separating on a magnet, the solution was transferred to a new tube. Then, another round of 6-8 cycles of PCR was performed to obtain sufficient molecules for sequencing. The cycle of PCR amplification was determined according to the amount of 1 ul amplified DNA, which was evaluated using FlashGel System (Lonza, 57063). Lastly, 300-500 bp DNA fragments were selected using Ampure XP beads with 0.5X volume plus 0.65X volume, then eluted in 15 μl ddH2O. DNA methylation libraries were sequenced on Hiseq2000 or Hiseq Xten platform (Illumina).

Low-input RNA-seq library preparation for zebrafish germ cells

For low-input RNA-seq libraries, we picked up 20-50 GFP-positive germ cells at each stage by micromanipulator and transferred samples into 7 ul DPBS with snap-frozen in liquid nitrogen immediately. Reverse transcription using REPLI-g WTA single cell Kit (QIAGEN 150063) as the kit's standard protocol. The amplified cDNA was purified using 1 volume Ampure XP beads, then fragmented to 200-400 bp by Covaris S2. Libraries were prepared with NEBNext Ultra II DNA Library Prep Kit (NEB E76455S) as described above and amplified with 6-7 PCR cycles. Sequencing were performed on Hiseq 2000 or Hiseq Xten platform (Illumina).

Histology

For juveniles at 35 dpf (approximately equal length of males and females around 1.7±0.1cm) and sex reversal adult fish, heads and tails were cut off. The middle body parts containing gonads were fixed in Bouin's solution (Sigma) overnight at 4°C. After dehydration, samples were embedded in paraffin and sectioned at 10 um thick. Hematoxylin and eosin staining (H&E staining) was then performed on the sections.

Morpholino injection

To analyze the function of *tet* genes, a knockdown experiment was conducted by the injection of a morpholino antisense oligonucleotide (MO), which was synthesized and obtained from Gene Tools.

*Tet* MOs including: *tet1* (AGATGCACTGTCTGAAGGGCTAATA) and *tet3* (TGGGACCCAATTTCCATCTGGCCTT). Approximately 4 ng and 8 ng (double injection, 4 ng each) MOs solution were injected in the one cell stages of *Tg* (*vasa::egfp*) transgenic embryos using a pressure micro injector. The phenotypes were observed under a stereoscope at different time points.

Counting PGC numbers in zebrafish germline

PGC numbers were counted using the squash method at the 5$^{th}$ day and the 9$^{th}$ day as described previously [12]. Briefly, for 5 dpf larva, we placed 5ul water on the slide, then put a larva in the water, pressed a coverslip onto the sample. For 9 dpf zebrafish, larva heads and tails were removed, then the body part containing gonads were dissociated with 0.25% trypsin and incubated at 30°C for 4 mins. Next, we transferred the body part on the slide, pressed a coverslip onto the sample. The number of PGCs was detected based on EGFP fluorescence under a fluorescent microscope.

*Tet3* CRISPER/Cas9 and genotyping

The *Tet3* mutant zebrafish was generated by using CRISPER/Cas9 as described [55]. *tet3* gRNA (ACCGAGTCGCACCCAAACTC) and Cas9 mRNA were synthesized by in vitro transcription, and then they were injected into 1-cell stage embryos. For CRISPER/Cas9 efficiency identification and genotyping. the 24 hpf embryos and caudal fin tissues were dissociated with 50 mM NaOH at 95°C for 10 mins (vortexes then repeat last process) and then balanced with equal volume 10mM Tris-Cl (pH8.0).

The primers used to PCR were *tet3* cas9 -S: AATAGCATGCCCAGGCTCAG and *tet3* cas9 –AS: GACAGTACAGAGCTCCTCAGG

Whole mount in situ hybridization

The 5 dpf larvae after the addition of PTU (Sigma-Aldrich P7629) were collected and fixed in 4% PFA/PBS overnight at 4°C. Next, they were dehydrated with MeOH and storage at -20°C. Larvae were rehydrated with DEPC-PBST and permeated with protease K (10 ug/ml) in DEPC-PBST. After permeation, larvae were washed with DEPC-PBST twice and then blocked in Hyb+ Buffer (50% Formamide, 5xSSCT, 0.5 mg/ml yeast RNA, 0.05 mg/ml heparin, 0.5% Tween) for 4 hours at 65°C. Next, larvae were incubated overnight at 65°C with DIG-labeled probe in Hyb+. After probes were removed next day, samples were washed with 25 mins 50% formamide/2xSSCT, 15 mins 2xSSCT, 30 mins 0.2xSSCT at 65°C. Samples were washed in MABT thrice at room temperature and blocked 1 hours in blocking buffer (2% Blocking reagent, 10% inactivated goat serum in MABT), and then were incubated with anti-DIG-AP Fab fragments in blocking buffer at 4°C overnight. Samples were washed in MABT for eight times at 30 mins, three times at 5 mins in staining buffer (100 mM Tris-Cl pH 9.5, 100 mM NaCl, 50 mM MgCl), then larvae were stained with BM Purple AP substrate in dark. Lastly, samples were fixed with 4% PFA/PBS for 30 mins and washed with PBST twice, then were steeped in 90% glycerol.

DNA methylation interference in zebrafish

(1) The experiment for 5-aza-2-deoxycytidine (5-Aza-dC) treatment at 9 dpf

5-Aza-dC power (Sigma-Aldrich (A3656) was dissolved in sterile deionized water (as stock solution 0.25 mg/ml). Then 5-aza-2-deoxycytidine (5-Aza-dC) treatment exposure was initiated at 9 dpf and terminated at 60 dpf in the vasa::egfp transgene zebrafish. 20-25 Juveniles were raised in a 1 liter tank from 9 dpf to 35 dpf, then transferred to 10 liter tank to 60 dpf, and finally transferred to zebrafish recirculation system. The fishes were fed twice per day: fairy shrimp mixed with 5-Aza-dC (0.005 mg/g, 0.010 mg/g and 0.015 mg/g respectively) in the morning, and normal diet in the afternoon. Half of the water within a tank was renewed per day. The fish genders were monitored under the fluoresced microscopy after drug treatment till 35 dpf and 60 dpf. The replicate details were shown in Table S5.

(2) The experiment for aromasin + 5-Aza-dC treatment during sex transition

Aromasin exemestano (*pfizer*) tablets were dissolved in sterile deionized water (as stock solutions of 25 mg/ml). The *vasa::egfp* transgenic adult females (3-4 months old) with normal mating behavior and fertility were randomly selected and allocated into three groups: the control group, the aromasin treatment-only group, and the aromasin+5-Aza-dC treatment group. 6-7 fishes were put into a 1 liter tank for each group. In the aromasin treatment-only group, the stock solution was mixed with fairy shrimp (1 mg/g) for feeding. In the aromasin+5-Aza-dC treatment group, 5-Aza-dC (0.035 mg/g) was added and mixed with fairy shrimp along with aromasin drug. Zebrafish were fed twice per day: drug food in the morning and normal diet in the afternoon. Half of the water within a tank was renewed per day

The ovarian fluorescent changes were monitored under the fluorescence microscopy weekly during the drug treatment period. After the continuous treatment for three months, the fishes were transferred to zebrafish recirculation system. At the stage of 4-5 months, the fishes were mated with normal females to check whether the fishes were successfully sex reversed and fertile or not. At the 3$^{rd}$ month and 5$^{th}$ month during drug treatment, we also counted and calculated the sex ratio for the control group, aromasin treatment-only group and aromasin+5-Aza-dC treatment group, respectively. Additionally, for each stage during sex transition, some of the gonads were collected for photographing and histological staining while the rest gonads were subjected to DNA methylation and RNA library construction. The treatment experiment replicate details were shown in Table S5.

Isolation and collection for zebrafish sex transition samples

The gonad was isolated from each individual fish during aromasin treatment and then separated from surrounding tissue and blood in D-hanks solution. The gonad divided into two part, one was immediately frozen at -80°C in RNA later (Sigma-Aldrich) to be used for the RNA sequencing and the other was in PBS for DNA extract.

For wild type sperm and sex reversal sperm, they were released from testis by gently pipetting in Hank's balanced salt solution (HBSS, Life) for 15 mins at 28°C. Then, the supernatant with swimming sperm was transferred to a new tube. The incubation-transfer was repeated 3 times and finally pelleted by centrifugation for 5 mins at 10000 xg. 100 ul Buffer X2 cell lysis solution (20 mM Tris-HCl pH 8.0, 20 mM EDTA, 200 mM NaCl, 80 mM DTT, 4% SDS and 250 ug/ml Proteinase K) was added to the samples, incubating at 55°C until the sample was dissolved (at least 1 hours) on a rocking platform.

Bisulfate-seq library preparation for zebrafish sex transition samples

Samples (including wild type/sex reversal sperm and gonads) were subjected to genomic DNA (>100 ng) extraction using by QIAamp DNA Mini Kit (QIAGEN 51304) following the manufacturer's protocol. Purified DNA spiked in with 0.5% un-methylated lambda DNA (Promega) was sonicated into 300 bp fragments with Covaris S2. Sheared DNA was transferred to a fresh PCR tube, NEBNext Ultra II DNA Library Prep Kit for Illumina (NEB, E7645S) was used for library construction according to manufacturer's instruction. DNA was end repaired and A-tailing by adding 7 ul NEBNext Ultra II End Prep Reaction Buffer and 3 ul NEBNext Ultra II End Prep Enzyme Mix. Samples were incubated in a thermal cycler at 20°C for 30 mins, 65°C for 30 mins, and finally cooled to 4°C. Adaptor ligation was performed by adding 30 ul NEB Next Ultra II Ligation Master Mix, 1 ul NEBNext Ligation Enhancer, 0.5 ul 200 mM ATP and 2.5 ul 25 mM cytosine methylated Illumina adapter. Sample were thoroughly mixed and incubated at 20°C for 25 mins. Adapter-ligated DNA was purified with 1 volume Ampure XP beads (Beckman Coulter A63881) to discard the short fragments and adapter-self ligations. Then bisulfite conversion was performed using the EZ DNA Methylation-Gold kit (Zymo Research) according to the instruction manual. Bisulfite-treated DNA was amplified using KAPA HiFi HotStar Uracil+ ReadyMix with 9-10 cycles. 300-500 bp DNA fragments were selected using Ampure XP beads with 0.5X volume plus 0.65X volume, then eluted in 15 μl ddH2O. DNA methylation libraries were sequenced on Hiseq2000 or Hiseq Xten platform (Illumina)

RNA-Seq for zebrafish gonads and gametes

Zymo Quick-RNA MicroPrep kit (Zymo Research, R1050) was used to extract RNA according to manufacturer’s protocol. RNA quality was assessed on agarose gel. The NEBNext Poly (A) mRNA Magnetic Isolation Module (NEB E7490) was used to isolate intact poly (A) + RNA from the 200-800 ng total RNA. Then mRNA reverse transcribed and amplified into cDNA and further library prepared using NEBNext Ultra II directional RNA Library Prep kit (NEB E7760) according to the manufacturer’s instructions. Libraries were pooled and sequenced in PE150 mode on Hiseq Xten platform (IIlumina).

DNA methylation data processing

Raw reads were trimmed to remove the adapters-containing and low-quality reads by Trimmomatic software with default parameters [56]. Clean reads were aligned by using Bismark(version 12.5) against to zebrafish genome assembly(Zv9) in paired-end mode with parameters -N 1 –X 600 [57]. The lambda genome was also included in the reference sequence as extra chromosome for bisulfite conversion rate calculation. After alignment, overlapped part of paired reads was clipped using clipOverlap function of bamUtil (https://genome.sph.umich.edu/wiki/BamUtil:_clipOverlap). PCR duplications were removed with Picard (http://broadinstitute.github.io/picard).

CpG methylation calls were extracted with Bismark methylation extractor function. Strands were merged to calculate the CpG methylation level per site. Average methylation level in each stage was the mean of methylation level at each CpG site. The CpG density was defined as the average number of CpG site per 100 bp.

Genomic elements were downloaded from UCSC Table Browser and quantification of average methylation levels of genomic elements was measured as the sum of the methylation level of CpG cytosine divided by the total number of CpG cytosine that reside in those genomic elements.

The R package (MethylKit) was used to cluster gonads samples hierarchically with "euclidean" distance by the "ward" clustering method [58].

Identification of DMCs and DMRs

To detect DMCs (Differentially Methylated Cytosines), 'mcomp' module from R package MOABS [59] was employed. CpGs with the cutoff of p-value was 0.01 and the difference of methylation level between two stages was higher than 0.2 were considered as DMCs. Candidate DMRs (Differentially Methylated Regions) were defined by a smoothing local likelihood method in R package bsseq [60]. DMRs containing at least 5 DMCs and whose difference level between two groups was higher than 0.2 were used for further analysis.

Identification of DMPs (Differentially Methylated Promoters)

The methylation level of each promoter was determined as the ratio of the number of alignments with C (methylated) over the sum of alignment with C and T for all CpGs in the promoter. DMPs were identified with two-tail Fisher's Exact Test and the p values were adjusted with Benjamini and Hochberg method. Promoters with adjusted p-value less than 0.01 and methylation level difference greater than 0.2 were considered as DMPs. DMPs clustering for zebrafish PGCs was performed using the 'pam' function of the 'fpc' package in R.

RNA-seq data processing

Reads with low-quality and adapters were trimmed by Trimmomatic software and then aligned by STAR (version2.5.2b) with default parameters [61]. Then filtering was performed to remove alignments with MAPQ < 20. The unique reads were used to calculate the fragments per kilobase of exon per million fragments mapped (FPKM) with Cufflinks (version 2.2.1) (http://cufflinks.cbcb.umd.edu). We filtered out genes with FPKM < 1 in all cell type. The gene expression of pluripotency and germline genes were computed by averaging from biological replicates. RNA-seq tracks for visualization were generated by bamCoverage tool in Deeptools2 with parameter "-normalizeUsingRPKM".

Time-course gene expression analysis

To analysis transient gene expression changes and to study the dynamics of their transcriptional activity from ovary to testis, we performed time course sequencing data analysis R package TC-seq (version 1.9） with default parameters. We selected genes with top 50% expression variance and membership value > 0.4 for further analysis.

Gene Ontology (GO) analysis

Gene ontology(GO) analysis of stage-specific gene expression was performed using DAVID [62]. Go terms with fisher exact test p value less than 0.05 were considered as statistically significant. Differential methylation promoter genes and difference time course cluster were analyzed using Metascape (http://metascape.org) [63] for functional enrichment.
